# DIscrete Dynamical Model of Mechanisms Determining the Relations of Biodiversity and Stability at Different Levels of Organization of Living Matter

**DOI:** 10.1101/161687

**Authors:** Yu. G Bespalov, K. V. Nosov, P. S. Kabalyants

**Affiliations:** VN Karazin Kharkiv National University, Ukraine

## Abstract

The paper aims at building the model of relations of biodiversity and stability at different levels of organization of living matter with the use of discrete dynamical models. The relations revealed in the study are illustrated by case studies of zooplankton community of the eutrophicated lake and the colorimetric parameters of the microalgae community of phytobenthos and phytoperiphyton. The results offer: (1) new approaches to estimating the risk of mass development of toxic cyanobacteria in eutrophicated reservoirs, and (2) new indicators for unmasking animals with protective coloring on the background of plant communities by image processing of digital photos.

## Introduction

Global climate changes increase the topicality of the problems related to the stability of communities of living organisms — inhabitants of our planet. The urgency of the problem of the stability of living systems in general also increases. At the same time, theapproaches to solving these problems, which are based on the notion of positive results of minimizing human impact on nature, are losing their former practical significance. This view corresponds toone of the two aspects of the strategy of harmonization of relations between human and nature, the essence of which in aphoristic form was expressed by B Commoner [1] as a "law" declaring that "Nature knows best."

Actually, another aspect of this "law" of Commoner becomes moreimportant, which supplements the warning against ill, conceived meddling in the mechanisms of performance of natural systems, by calling for comprehension of these mechanisms.

Regarding this second aspect, it should be necessary to mention the concepts linking the stability of communities of living beings and natural systems in a whole, with biodiversity The opinion on the positive role, in this sense, of biodiversity has become widespread, but at this time it does not have an unambiguous confirmation based on studies of actual data[2-5].

The ambiguity of information on the role of biodiversity in maintaining the stability of living systems makes us recall the opinion expressed by R. Margalef [6] that "the ecologist sees in any measure of diversity the expression of the possibility to construct a system with feedback".

Now it’s time to note here that the complexity of ecological systems in many cases makes it difficult to collect an initial data in the volume necessary for constructing feedback systems based on the approach involving description of the entire set of causal relationships between the components of the system, which determine the nature of system’s homeostasis, including its absence.

The original class of mathematical models for formalized description of dynamical systems of different character [7-14] was developed with authors’ participation These models, called discrete models of dynamic systems(DMDS), allow the researcher to reduce the requirements for the initial actual data in the process of building the system’s model. To be exact, this class of models enables to construct the models with the use of relatively small observation tables, having gaps, and observations with disturbed time order. In other words, for model’s identification, the initial unordered data can be used, in contrast to various dynamical models like time series. The initial data may present levels or grades of certain property, that reduces demands to the quality of measurement.

An identified model shows, in the matrix or graph form, the structure of between, component and intra, component relationships of the system The relationships express the presence/absence of positive or negative between, and intra, component influences, which can be described by the set)(0,0),(+,0),(0,+),(-,0),(0,-),(-,+),(+,-),(-,-),(+,+). The component subject to a positive influence tends to increase its value 2amount of property, characteristic3 in the following time; correspondingly, a negative influence forces the component to decrease its value So, the dynamics of the system is determined by the structure of relationships and the current 2present3 state of the system For mathematical details, refer [8-10].

On the basis of the relationships structure obtained in such a manner, for a given initial conditions, the idealized trajectory of the system (ITS)can be calculated The ITS is actuality a cycle of changes of components’ values Integer scores(beginning with 1)are used as the values of the components in the system The ITS makes it possible to analyze the dynamics of values of Shannon’s and other indicators of diversity and evenness proposed in ecology [6,15-17].

The ability to analyze the values of these indicators in dynamics offers the challenge of eliminating the problems related to the difference between these values in different periods of performing the communities of living beings The expression of values in conditional scores, in a number of cases, enables to eliminate biologists’ reasons regarding the use of the Shannon index and other indicators of diversity and evenness calculated on the basis of data on abundance, biomass and energy flows in trophic chains The essence of these objections is that these indicators do not reflect the role of rare species in performance of biological communities.

This work aims at studying the abilities provided by the DMDS models in development of new approaches to the investigation of the role of diversity in performance of communities of living organisms.

## Material and methods

For purposes of the study, the aspects of diversity and evenness that may have relation to the nature of the sets of strategies for maintaining the stability of a living system was analyzed Such analysis was performed for the ITSs built for biological systems that preserve their stability to a different extent The zooplankton communities of Lake Sevan(Armenia)at different stages of its eutrophication are considered as such systems The initial data of observation on zooplankton at these stages were taken from [18].

Also, the ITSs based on the distribution of brightness in spectral bands of reflection of plant pigments on photo images of the cyanobacterial community was analyzed The photo of the cyanobacteria community is downloaded from the public website of space images (https://lance.modaps.eosdis.nasa.gov) To determine the brightness in spectral bands of reflection of plant pigments, the relations of the RGB-model components were analyzed, which correspond to these bands(with a certain degree of roughening).

The working hypotheses suggested during the analysis of ITSs were tested by statistical methods The images obtained by digital processing allowed us to confirm the correctness of assumptions

For calculation of the ITSs, the measure of proximity with the use of Spearman correlation and the approach on the base of Liebig law of minimum [8] were applied.

## Statement of the problem and discussion

Stating the problem, we should choose a sufficiently productive understanding of the meaning of the concept of diversity. The understanding of biodiversity as a diversity of strategies for maintaining the stability of a living system is, in our view, quite reasonable. The use of species diversity and other types of biodiversity increases informativeness of this diversity in the context of such understanding. We have in mind the informativeness only in the context of the problem of relations between biodiversity and stability of living systems; in the scope of this study, other aspects of biodiversity(ethical, aesthetic, world, outlook related etc)are not considered Besides, in the present study the stability of biological systems caused exclusively by the mechanisms of homeostasis, that provide self, regulation, is under considered Such mechanisms provide the ability of an open system to maintain the stability of its internal states through coordinated reactions aimed at maintaining a dynamic equilibrium.

This note is necessary in connection with the following reasons There are cases when the stability of biological systems is provided, roughly speaking, by the law of large numbers and factors that minimize the intensity of bioproduction processes A well-known example of such a kind is peat bogs accumulating huge amounts of organic matter and biogenic elements The involvement of these biogenic elements in the cycle of bioproduction processes is minimized by the conservation effect of humic substances.

The stability of biological systems provided by the mechanisms of homeostasis is a sort of dynamic equilibrium, which assumes fluctuations in the values of system’s parameters within in a certain cycle.

The discrete dynamical model of such mechanisms makes it possible to represent this cycle in the form of ITSs providing the parameters’ values at different moments of the cycle These values are expressed in(conditional)scores, which makes it possible, to a certain extent, to ignore the random fluctuations of the parameters’ values within the limits of the scores’ values It should be noted that the ITS is an *idealized* trajectory, which represents the cycle caused exclusively by the structure of relations between the components of the systems This structure is described in the framework of the DMDS model and enables to calculate the ITS based on given initial conditions.

If the structure of inter-component and intra, component relations of the model is changed, we, in fact, obtain the model of another system.

A change in the system accompanied by violation of the character of its homeostasis can occur as a result of external influences. Such influences correspond to changes in the set of values of the system’s components. As a result, in this ITS, the states that previously not were observed can appear Such new combinations can arise at one or several moments of the ITS. The probability of arising new states decreases with the increase of the trajectory’s length and unique states corresponding to the number of moments.

In this regard, the number of unique states(ie, trajectory’s length)can be considered as an index of diversity related to the stability of the system and the ability of system’s homeostasis mechanisms to resist to external influences. We do not tell on the diversity of the components of the system(for example, members of the community of biological specie), as it remains constant in the considered model In fact, we have in mind the diversity of states(which may be interpreted as a variety of strategies used by the system to self-preservation).

There is also a certain possibility that increase in the ITS length will increase not only the number of states but also a range of values of indices of evenness of these values Such the possibility increases with the growth of a degree of mutual distance between maximum and minimum values of different components in the ITS

For determining the degree of evenness, the Shannon index is often used [6, 15,16]. Likely, in a number of cases, simpler expressions can be used for estimation of evenness of the values of a pair of components. As diagnostic characters of the state of homeostasis mechanisms, indices of range’s values of these characters can also be used

Taking this into account, it makes sense to state the problem of determining the state of homeostasis mechanisms with the use of the following procedures

The initial procedure:constructing ITSs of systems with different mechanisms with the help of the DMDS on the base of actual data. The difference between these mechanisms also determines the difference in the systems themselves. Systems can differ in the efficacy of homeostasis mechanisms and ability to maintain their stability. The zooplankton community ofa eutrophicated lake in the states described below can serve as an example.

1st state:the nature of dynamic equilibrium corresponds to the conditions in which there is a very small chance of mass development of toxic cyanobacteria in the lake This condition was recorded for a long period with a relatively weak anthropogenic eutrophication of Lake Sevan from 1937 to 1957.

2nd state:the nature of dynamic equilibrium inherent to the 1st state is disturbed, and mass development of cyanobacteria is possible in the near future This state corresponds to a relatively short, transitional period from the 1st to the 3d stage of Lake Sevan’s eutrophication from 1958 to 1961.

3d state:the nature of dynamic equilibrium, for which the possibility of mass development of cyanobacteria was realized and being constantly observed, has been established This corresponds to the eutrophication period of Sevan from 1962 to 1969.

Hereinafter, we will talk about the 1st, 2nd and 3d states of Sevan zooplankton and, respectively, the 1sd, 2nd and 3d periods of anthropogenic eutrophication of Lake Sevan.

As already mentioned, the 2nd state is transitional from the 1st state to the 3d, and the 2nd period is transitional from the 1st period to the 3d one.

We study the characteristic features of the cycle of the system when system’s homeostasis is maintained These features enable to distinguish the system to be identified from other studied objects, parameters of which don’t have a systemic nature and should not change over a certain cycle related to the homeostasis’ mechanisms. Grass canopy’s communities and locust populations that are barely visible on the canopy’s background due to protective coloring may serve as an example of such a system and such a non-systemic object [13]. Another example, mentioned below, is the community of microalgae on the bottom of a pond and a fish (pike) on its background.

Later, the type of ITSs was analyzed. The aim of the analysis is to find systemic parameters that reflect the essential features of the relationships structure and the dynamics of the components of the system under study. These features should distinguish this system from others ones or from objects that do not have such a systemic nature. These system parameters can be used to identify certain states, using statistical methods, or by determining the location of certain objects on the background of some biosystems with the help of image processing.

In this paper, the following results were obtained using such procedures

The first example is a formalized description, in the form of ITS, of some aspects of the dynamics of the zooplankton community of Lake Sevan for different periods of its eutrophication, the analysis of the ITSs, developing and testing the working hypotheses based on this analysis.

For these three periods, the ITSs describing the dynamics of the number of four species of zooplankton *Keratella quadrata, Filinia longiseta, Daphnia longispina*, and *Cyclops strennus* were constructed [8]. The ITSs are shown in Fig. 1, 2, 3.

**Fig 1.**
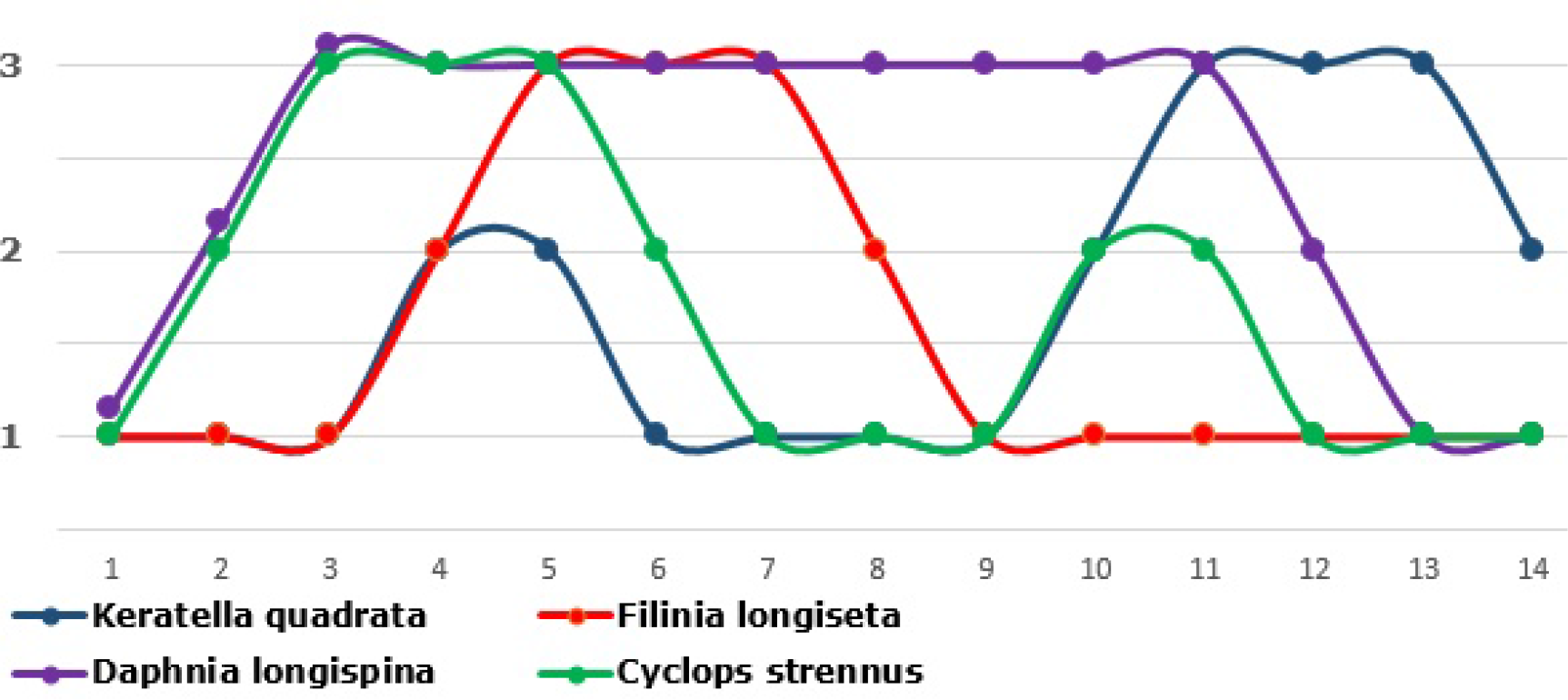
Idealized trajectory of the system, reflecting the dynamics of the numbers of zooplankton species in the 1st period of anthropogenic eutrophication of Lake Sevan The number of species takes 3 levels 21 1 low, 2 1 medium, 3 1 high3, the trajectory’s length is equal to 14 moments Any moment defines a state of the system comprising the number of 4 zooplankton species For better visual presentation, the trajectory’s lines are smoothed.

**Fig 2.**
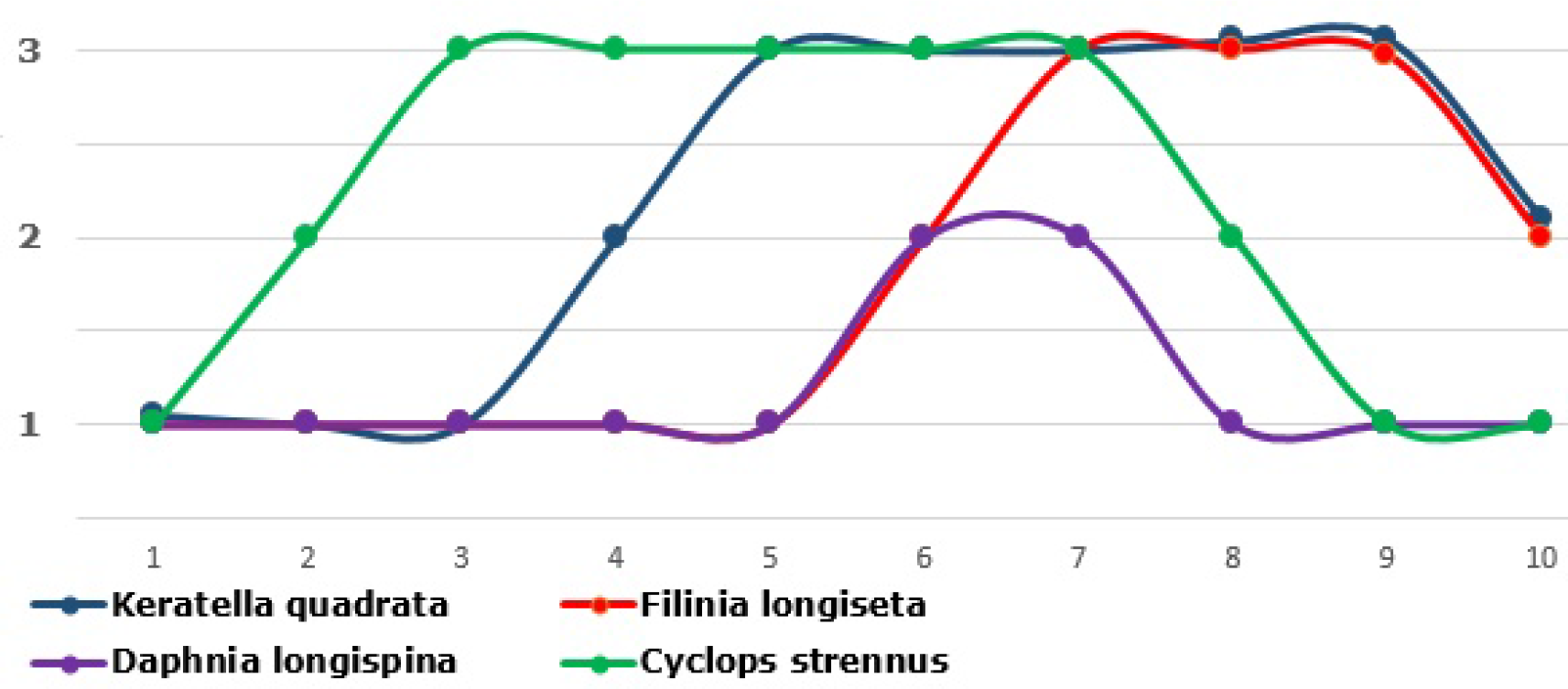
Idealized trajectory of the system of the number of 4 zooplankton species in the 2nd period of anthropogenic eutrophication of Lake Sevan.

**Fig 3.**
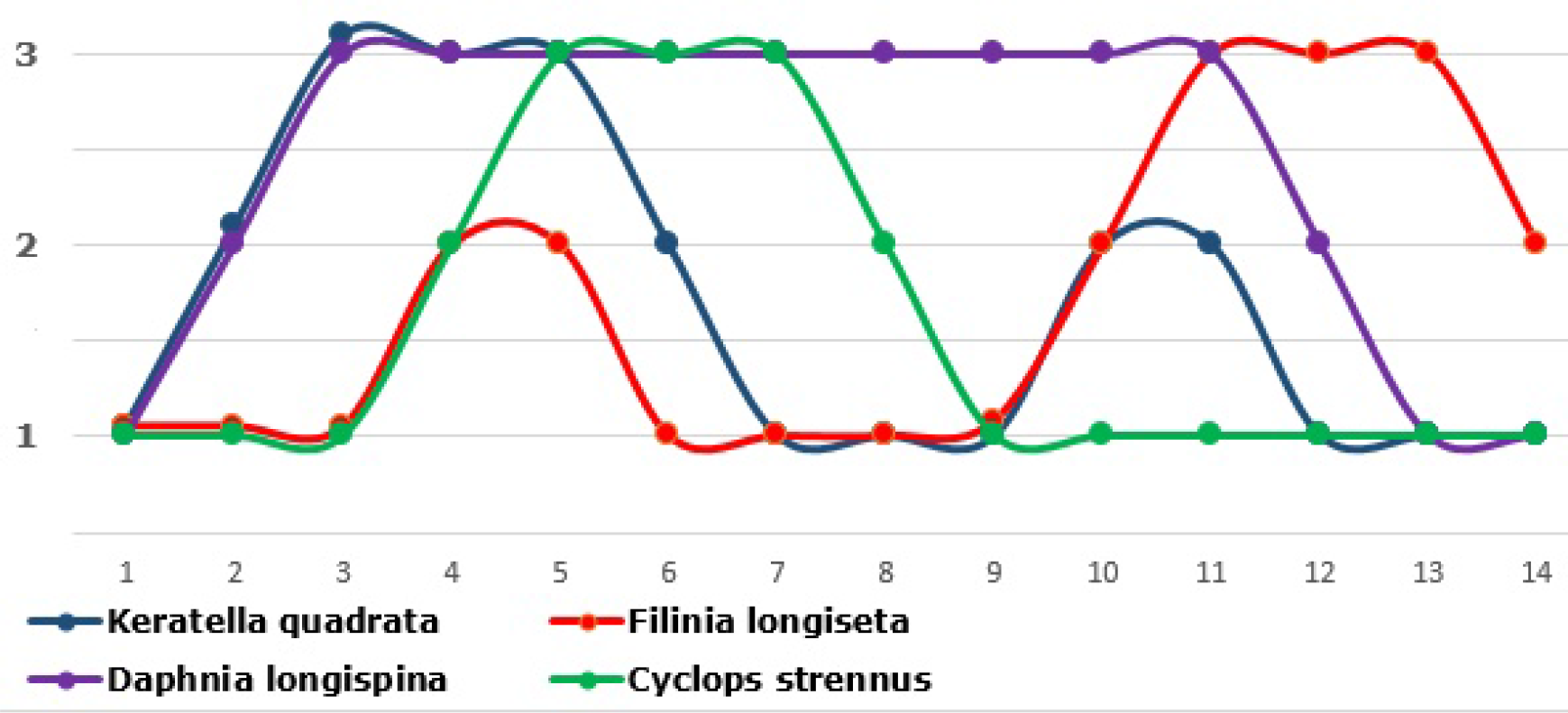
Idealized trajectory of the system of the number of 4 zooplankton species in the 2nd period of anthropogenic eutrophication of Lake Sevan.

From the ITSs in Fig 1, 2, 3, it follows that the 2nd period cycle’s length is the shortest(10 moments)in comparison to the length of the 1st and 3d period(14 moments each). This corresponds to a smaller number of non-repeated states of the system In accordance to said above, a smaller number of these states should correspond to a lower ability of the system to maintain its stability.

The real data confirm this. The second state of the Sevan ecosystem was short, lived and transitional between the first and third one, which were differed in such significant parameter as massive development of toxic cyanobacteria. Another feature of instability for the second state, marked earlier in the second period, is the statistically significant correlation between the frequency of magnetic storms and the number and biomass of *Daphnia longispina* [8]. In the first and third period, there was no such the correlation This reason allows us to consider the correlation as an additional criterion for recognition of the second period.

Comparison of the ITSs in Fig 1, 2, 3 enables to distinguish the differences between the 2nd state, on the one hand, and the 1st and 3d states, on the other, in the distance of maximum values of the number for different species. In the ITSs for the 1st and 3d periods, the set of moments with maximum values of *Keratella quadrata* and *Filinia longiseta* are far apart, but for the 2nd period, they almost coincide To a lesser degree, similar differences are peculiar to the number of *Keratella quadrata* and *Cyclops strennus*.

There are some reasons to assume, that a greater degree of coincidence of maximums for species’ numbers will accompany by a greater degree of evenness of these values. Such a result will be obtained by determination of the degree of evenness with the use of Shannon index when the modulus of the difference between two values of characteristics is used. The degree of evenness for the values of K*eratella quadrata, Filinia longiseta, Keratella quadrata* and *Cyclops strennus* is higher for the 2nd period|—compared to the 1st and 3rd ones. This contradicts arguments said above regarding a less stability of the 2nd period in the context of a widely-held(but criticized)view on the relations between evenness and stability. At the same time, as mentioned, the analysis of the ITSs for the three periods of Sevan eutrophication is consistent with the concept of relations between stability and diversity.

Here, we are dealing with the diversity of strategies of performance of the system(which the ITS’s length corresponds to) and the stability, indicator of which, in the case of the aquatic ecosystem under investigation, may be a relatively long-term existence of the state with frequent outbreaks of biomass of cyanobacteria or without such outbreaks.

It seems reasonable(without tight relations to one or other concepts)to use such indicators of diversity and evenness, which measurement enables to identify certain ecosystems’conditions. In this example, the indicators enabling identification of the state of an aquatic ecosystem that constitutes, in the near future, a thread of mass development of toxic cyanobacteria The thread of such a state is very high nowadays, for example, for Lake Kinneret, which is the main source of potable water for Israel [19,22]. For the anthropogenic eutrophication of Lake Sevan, this dangerous state is the 2nd state mentioned above. Its features indicated the rapid rise of frequent outbreaks of biomass of toxic cyanobacteria in the ecosystem of the lake.

The analysis of the ITSs in Fig 1, 2, 3 can assist to formulate working hypotheses concerning systemic parameters enabling to identify such a state in advance in statistically significant manner. In particular, indicators of evenness of the number of species can serve as a diagnostic character of this state. Such indicators calculated by the numbers of the above, mentioned species will not be very important because the species composition in eutrophicated lakes plays an important role But such an approach with the determination of the number of larger taxonomic or other groups of zooplankton can be useful for estimation of the risk of mass development of toxic cyanobacteria with the emergence of threats to biosafety of potable water consumption.

In the study, the working hypothesis based on the distance between maximums of the number of species *Keratella quadrata* and *Filinia longiseta* in the trajectory were tested, for the 1st and 2nd periods. According to this hypothesis, the first and second periods should significantly differ by variances of the modulus of the difference between the number of *Keratella quadrata* and *Filinia longiseta*.

Initial data of the number of *Keratella quadrata* and *Filinia longiseta* were normalized by the means of samples of these species correspondingly. The variances of normalized samples significantly differed(at the level of 001) according to the Fisher *F*-test.

The second case study presents the results of formalized description of some aspects of the dynamics of colorimetric parameters in the community of cyanobacteria. This community in an ecological sense is similar to many communities of microalgae that form clumps on the surface of water or various substrates We use colorimetric parameters, which in a certain sense reflect the quantities and relations of various plant pigments and can be obtained by computer analysis of components of the RGB digital model on digitalimages. The ITS in Fig 4 represents the dynamics of the following colorimetric parameters:

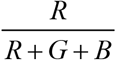 – reflects the amount of orange, yellow pigments dominated in old, dying and dead plant cells;

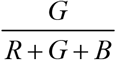 – reflects the amount of green pigment of chlorophyll dominated inyoung, actively photosynthetic cells;

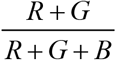 – reflects the total number of all cells (young and old, dying and dead);

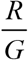 – reflects the value of the "yellow, green index", which is an indicator of pigment diversity [23]

The dynamics of these colorimetric parameters in Fig 4 shows a marked difference in the colorimetric parameters 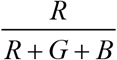 and 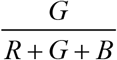, as well as the even more significant difference of maximums of 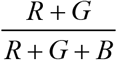 and 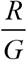. Due to the greater difference, the relation of this pair of colorimetric parameters is more suitable for usage in image processing for specific purposes This reason will be used for image processing of some biological objects on the background of microalgae communities. It is assumed, that the relations’ structure of the colorimetric parameters of these communities corresponds to the dynamics of changes similar to those presented in Fig 4.

**Fig 4.**
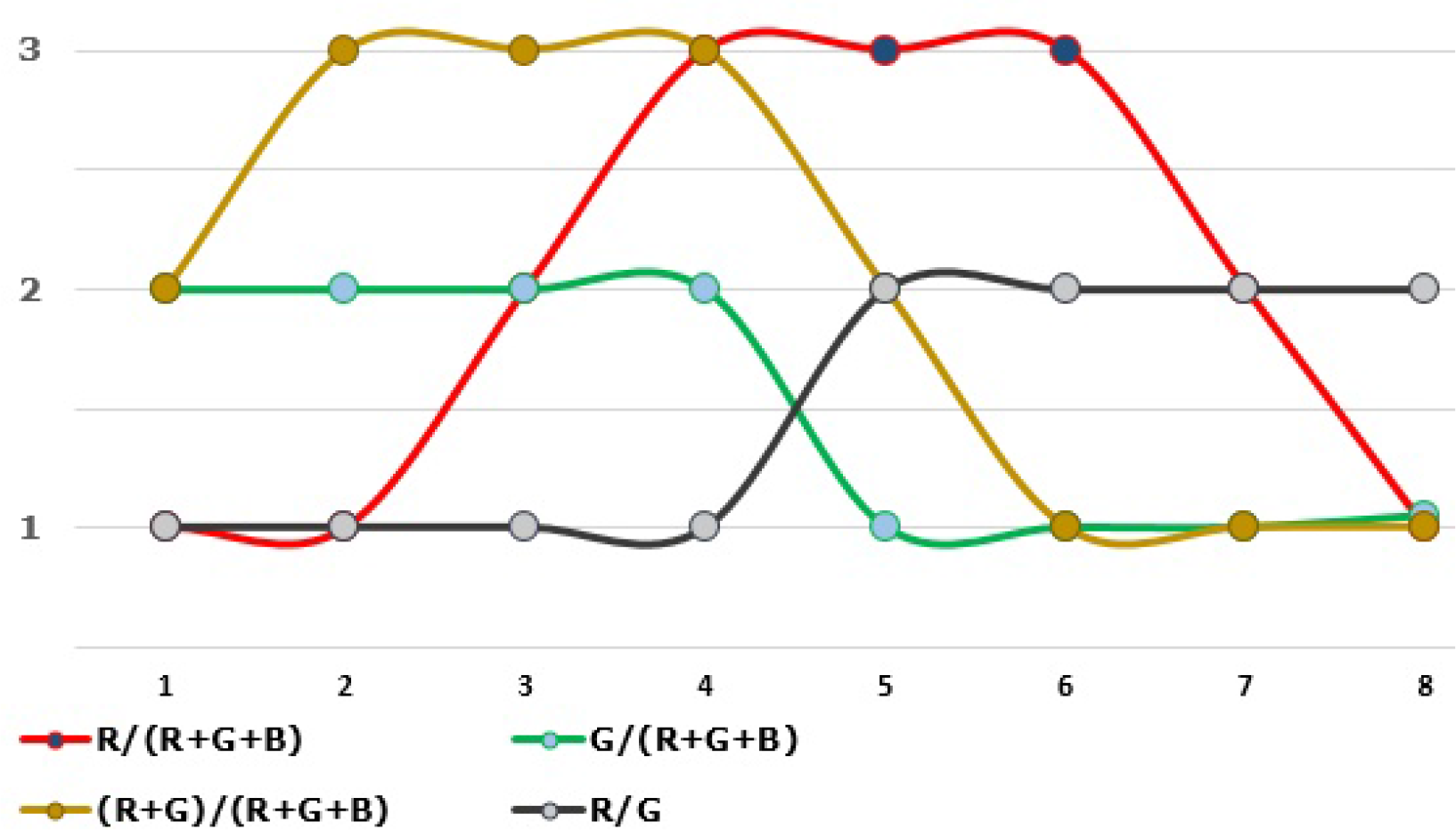
Idealized trajectory of the system reflecting the dynamics of colorimetric parameters of the community of cyanobacteria

The possibility that this is not true is quite high for the biological objects, which photos taken on the background of microalgae communities, in particular, for animals with protective coloring. Accordingly, it is reasonable to expect a large difference in the diversity of combinations of the specified colorimetric parameter. So, these features will unmask an animal on the background of thisplant community.

In a number of cases, the use of the DMDS enables to reconstruct the dynamics of the colorimetric parameters of plant communities on the basis of an image taken at a single moment. This technique has the working name of *rechronization*. This method is based on the assumption that different parts of a plant community change their colorimetric parameters according to a certain cycle, but are at different moments of the cycle at a moment.

The ITS shown in Fig 4 is obtained with the help of rechronization The trajectory demonstrates above-mentioned relations between the parameters 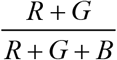 and 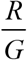. The nature of these relations implies a high probability of coincidence of high values of one of these parameters with low values of another. In turn, from this, it follows a sufficiently high possibility of relatively low diversity in the values of the product of these two parameters. So, the following working hypothesis can be suggested:the product 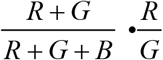 can serve as an indicator of a certain unmasking effect The effect unmasks an animal with protective coloring on the background of the community of microalgae biofilm.

Fig. 5 confirms this hypothesis This image was obtained as follows. A digital noise(to simulate unfavorable conditions of photo-taking) was added to the initial image of the fish (pike). The noised image was processed using the introduced indicator in the following manner. The noised image was split into segments, and each segment, in turn, was divided into smaller segments(sub, segments). For each first-order, large segment the following sample was generated:the value 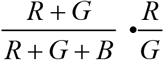 for each sub, segment constituted a case. Then, for each first-order segment, the value of the variance of its sample was calculated, and this value was used for visualization in Fig.5c.

**Fig 5.**
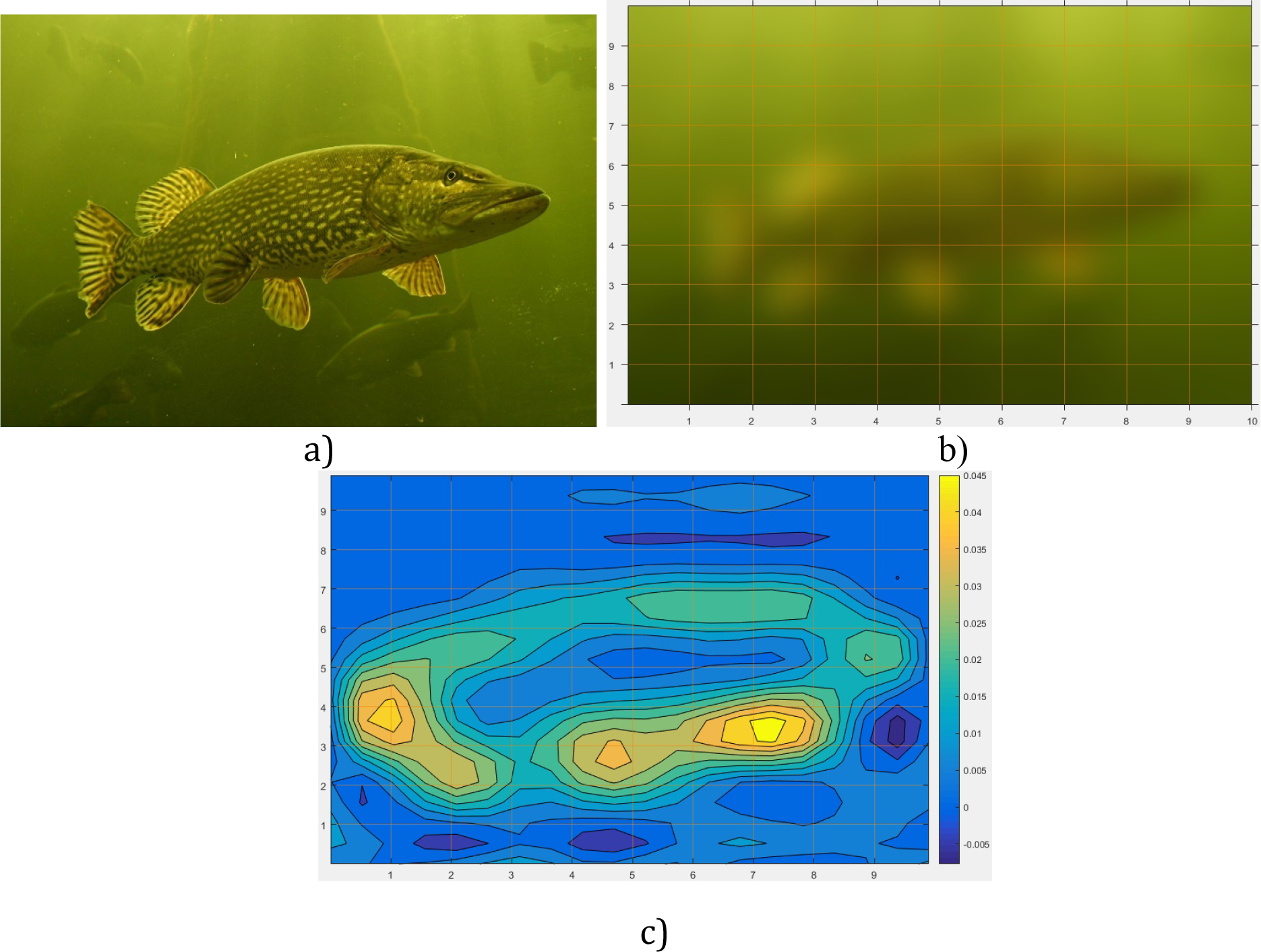
Result of image processing of the fish on the background of microalgae film, a) original image, b) noised image, c) processed image

The contour of the fish obtained by such image processing shows, that the value of the variance is higher for the contour in comparison with the background, that is consistent with the working hypothesis. Though this contour in the noised image is less informative than in the original image, this approach may be useful for development of remote registration methods of fishes, that play a significant role in many aquatic ecosystems.

## Conclusion

In the paper, the abilities for development of new approaches to the use of the DMDS model for studying the role of diversity in performance of communities of living organisms were shown.

These approaches offer the challenge for development of new methods of applied ecology. They may include, in particular, methods for predicting the risk of mass development of toxic cyanobacteria in water reservoirs, as well as remote methods of studying animals with protective coloring.

The resemblance in using the DMDS models as a tool for suggesting working hypotheses with certain aspects of the "algorithms of invention", proposed in his time by GS Altshuller, is also of some interest.

